# Decoupling badger and sett distributions for improved bovine tuberculosis management

**DOI:** 10.1101/2025.11.10.687598

**Authors:** Virginia Morera-Pujol, Andrew W. Byrne, Damien Barrett, Philip Breslin, Guy McGrath, David Quinn, Simone Ciuti

**Affiliations:** School of biology and environmental sciences, University College Dublin, Belfield, Dublin 4, Ireland; Department of Agriculture, Food and the Marine (DAFM), Dublin 2, Ireland; Centre for Veterinary Epidemiology and Risk Analysis, School of Veterinary Medicine, University College Dublin, Belfield, Dublin, Ireland

## Abstract

Bovine tuberculosis (bTB), a zoonotic disease caused by *Mycobacterium bovis*, continues to challenge eradication efforts in Ireland and the UK, partly due to the role of the European badger (*Meles meles*) as a wildlife reservoir. Traditional management strategies often rely on sett (burrow) locations to infer badger distribution, which implicitly assumes a correlation with abundance. This study uses data from Ireland’s national badger culling and vaccination programme (2019–2025) to decouple badger and sett distributions using spatial point process modelling via log-Gaussian Cox processes. By separately modelling the environmental drivers of main sett and badger distributions, and validating outputs for ecological realism with independent badger body weight data, we demonstrate that sett and badger densities are governed by distinct ecological processes. Sett densities are driven by landscape features such as elevation, slope, and proximity to forest edges, while badger densities are more influenced by recent culling history and pasture availability. Our results reveal a spatial mismatch between high-density sett areas and high-density badger areas, highlighting the need for refined metrics in wildlife-based bTB management. These findings underscore the importance of integrating independently derived wildlife distribution models into disease control policies for more sustainable and effective bTB management.

## Introduction

The increasing anthropisation of natural habitats is accompanied by challenges for the conservation and management of wildlife, like habitat fragmentation and loss, and population declines in many species unable to cope with the fast-changing environment (Parmesan 2006, Dobson et al. 2021). One additional and sometimes overlooked consequence of the human encroachment in natural and semi-natural ecosystems is the increased risk of zoonotic diseases, as many have wildlife vectors and reservoirs (Marrana 2022, Galindo-Gonzalez 2024).

An example of a zoonotic disease with wildlife reservoirs and significant implications in human-wildlife coexistence is bovine tuberculosis (bTB), caused by *Mycobacterium bovis.* This disease affects cattle but can also be transmitted to humans, having been the cause of an estimated 12,500 human deaths worldwide in 2016 (WHO et al. 2017). Within Europe, most countries have managed to achieve officially TB-free status – or at least achieve insignificant bTB levels, low enough to meet the official freedom threshold (<0.1% confirmed herd prevalence) – with mainly just cattle management measures, however it is still an issue of high concern in countries such as Ireland and the UK (Broughan et al. 2016, More et al. 2018). In Ireland in particular, herd incidence was over 5% between June 2023 and 2024, leading to the removal of 32,677 reactor (e.g. positive to a bTB test) cattle (Department of Agriculture 2024).

There are many societal and environmental factors interfering with bTB eradication in Ireland and the UK, but one of the causes that is generally accepted for both countries is the role of the European badger (*Meles meles,* badger hereafter) in the transmission of bTB (Bourne et al. 2000, Byrne et al. 2013b). However, badgers also play an important role in the ecosystem, and therefore were given protection in Ireland in 1976, which makes their management and conservation a complicated issue (Book 2024)

Badgers are a social species, with complex population dynamics and distribution that depend on the spatial structure of territories and the social group size (Rogers et al. 1996). Habitat structure drives the distribution of territories, with more productive habitats being able to support more territories and therefore more social groups, while more granular spatiotemporal variations in food availability drive group size (Byrne et al. 2015). Through a density-dependent process, badgers living in larger groups tend to have lower body weight, and consequently a worse body condition that has been linked to a higher susceptibility to parasites and other infections (Macdonald et al. 2002). However, some studies show how their social habits might complicate the issue of bTB transmission, as the increased sociality of larger groups might facilitate behaviours that help avoid habitats where infection risk is high (Albery et al. 2020), and as the vaccination of some badgers in a social group or area can provide protection against the disease to unvaccinated badgers of the same group (Carter et al. 2012).

The first bTB eradication actions in Ireland (1950s to 1980) focused mainly on herd management measures (More and Good 2006). In the late 1980s and early 2000s, two different experiments tested the effect of the removal of badgers, providing evidence to suggest that badger culling helped reduce bTB risk in cattle in high bTB prevalence areas (Máirtín et al. 1998, Griffin et al. 2005). Since 2004 a nationally coordinated culling programme has been in place by which badgers up to 2km of a breakdown farm are removed, under the assumption that reducing the density of the host species would reduce contact and transmission rates within and between species (O’Keeffe 2006). Setts are revisited at least once a year once they enter the system, and if signs of activity are detected, another capture event will be triggered (Byrne et al. 2013b).

In order to design a more sustainable badger management programme, field trials to investigate the efficacy of BCG (Bacillus Calmette-Guérin) vaccination in badgers (Aznar et al. 2011, Gormley et al. 2017) and effectiveness of badger vaccination relative to culling in cattle (Martin et al. 2020) were undertaken. The former demonstrated significant benefits in reducing incidence of bTB in vaccinated badgers, relative to placebo treated badgers, while the latter suggested that vaccination was not less efficient than culling at controlling bTB spread in cattle. On the basis of these pieces of evidence, a vaccination programme began in 2019 in selected areas in the country and has been expanding since then. By 2021, 20,000 km^2^ were designated as vaccination areas in Ireland (Ryan et al. 2023). Despite the expansion of the vaccination programme, it is still unclear what percentage of the population has been, or would need to be, vaccinated for the effect to be sustained in time, and what success will vaccination have in curtailing bTB expansion in areas of high badger density (Byrne et al. 2013a, Martin et al. 2020).

Here we set out to use the data generated by both the culling and vaccination programmes coordinated by the Department of Agriculture, Food, and the Marine (DAFM) of the Irish Government to model the distribution of badgers in Ireland. Using data on the location of both setts and captured badgers, and through point pattern models, we aim to (1) identify environmental factors that affect the distribution of both setts and badgers in the Irish landscape and (2) model the relative abundance of setts and badgers separately, assessing their differences, if any.

Badgers are expected to have lower body weights when living in larger groups (Macdonald et al. 2002), while it is also known that larger territories (and therefore lower sett density) are found in areas with less availability of resources and therefore where we expect to find badgers of lower body weight (Byrne et al. 2015). Therefore, we exploit these well-known relationships to (3) test the ecological realism of our sett and badger density models with independent data on body weight collected within the vaccination programme. When using our predicted badger and sett density distributions to explain spatial structure in badger weight, we expect badgers captured in areas with lower sett density to weigh less, and we expect estimated group size to have a negative relationship to body weight.

## Methods

### Wildlife data

#### bTB eradication scheme in Ireland

Information on the location of the setts was obtained from the national-scale badger management programme ran by the bTB eradication unit at the Department of Agriculture, Food and the Marine (DAFM) from the government of Ireland, which runs both a culling and a vaccination programme as described above. The main differences between the two programmes are in the animal intervention (vaccinated animals may be recaptured which needs to be taken into account), and in the duration of the fieldwork bouts (capture events from now on) which last six days in the vaccination programme and 11 days in the culling programme.

#### Setts

Setts are large burrow systems with several entrances and can be divided into main and non-main setts. Typically, there is only one main sett per social group, and therefore main setts can be used as a proxy for the presence of one social group (Etherington et al. 2009). Although setts entered the database when they were found as a response to a bTB breakout in a farm, once they were found they were visited relatively regularly to check for signs of activity, regardless of whether they were in a vaccination or a culling area. All setts where one or more badgers have been captured are visited at least once per year and scored for field signs of activity (Martin et al. 2017). From the original data we obtained the sett ID number, the date of the last field visit, the ID of the capture event, and whether it was a main sett or not. For our analysis we filtered the data to retain only setts last visited between from 2019 and May 2025 (as collapsed or disappeared setts stop being visited) and classified as main (as main setts can be generally assigned exclusively to one social group and therefore can serve as proxy for them).

#### Badgers

Information on the location of captured badgers was obtained from the same wildlife management programme, but in two different datasets pertaining to the culling and the vaccination programmes. As with the sett data, badger data were filtered to retain only captures between 2019 and May 2025, and the recaptures were removed from the vaccination dataset, so every animal only appeared once (at first capture). Badger capturing effort in Ireland is strongly concentrated at sett entrances or as close by as practicable, in the case of the presence of density scrub, on paths/ “runs” (Martin et al. 2017), and therefore the sett coordinates are assigned to the capture. In the minority of cases where badgers are not captured near setts, they were assigned by field staff the coordinates of the geographically nearest sett. Badger locations were then jittered randomly in a 500m radius around the sett location, as point process models cannot have duplicated points and usually more than one badger were assigned to each sett. Additionally, a thinning procedure was applied using the *spThin* package (Aiello-Lammens et al. 2015), where observations were removed if they were within 250 m of another location. This is a requirement of the modelling approach, as integration tends to fail if points are too clustered, and resulted in the loss of only 6.9% of the observations. Badger weight was recorded for all vaccinated badgers in the dataset before the cleaning and thinning process, in kg.

#### Effort

For management purposes, the country of Ireland is divided in 2 by 1.5 km tiles called “quartiles”. As these are the units used by the teams performing the culling and vaccination efforts to identify the location of the setts, we used them as our base unit to calculate the effort as well. Since the duration of the capture events were different for vaccination (six days) and culling (11 days) programmes, the effort was calculated as number of days spent on each quartile, instead of number of capture events. Therefore, a quartile that had been host to 5 culling capture events during the study period would have an effort value of 55, whereas a quartile that had hosted 5 vaccination capture events would have an effort value of 30.

Since setts are a (relatively) permanent fixture in the environment, there is a limitation on the number of setts that can be found in a quartile, that is, once all setts have been found no more will appear. To reflect this, we log-transformed the effort before using it for the sett model, but we used the untransformed effort for the badger model, as setts can be recolonised after being emptied by the culling programme, and therefore the number of badgers found in a quartile does not necessarily saturate. The specific way in which these efforts are included in the model is explained below in the modelling section.

### Environmental data

As the models used for this study are computationally expensive, it is not recommended to perform stepwise methods to select relevant covariates. Therefore, we selected the covariates to use for the model based on prior knowledge of environmental variables that affect badger distribution, and we used the same environmental variables for the setts and badgers model, to be able to compare how the two different processes are affected by the different environmental variables.

Elevation and slope are known to affect badger distribution in Ireland (Byrne et al. 2014). Therefore, we used the Digital Elevation Model (DEM in m) over Europe at a 100m resolution from the Eurostat database. The elevation raster was cropped to Ireland and rescaled to a 1km resolution to have all covariates at the same resolution. The slope was obtained from the elevation layer using the *terrain* function from the package *terra* (Hijmans 2025) according to (Horn 1981) and expressed in radians.

We were particularly interested in whether the badgers were attracted to grasslands and “enriched grasslands” (pastures) for the implications this might have in the transmission of bTb to cattle (Murphy et al. 2022). To assess this, we downloaded the CORINE land use vector layer from the European Union’s Copernicus Land Monitoring Service. From these, we selected the polygons representing natural grasslands and pastures, and then calculated, over a 1km^2^ grid, what percentage of each grid cell was covered by either natural grasslands or pastures. This way, we obtained a continuous surface indicating the % of coverage of pastures and grasslands in a 1km^2^ grid.

Badgers have been positively associated with forested habitats in Ireland (Byrne et al. 2014, Elliott et al. 2015), with anecdotal evidence suggesting that they might be using forest edges rather than living in fully forested areas. Therefore, from the same CORINE land use vector layer we selected the polygons representing coniferous, mixed, and broadleaved forest and calculated the distance, in meters, to the edge of these polygons. We used positive values to express distance to the forest edge from areas outside of forests, and negative values to express distance to the forest edge from areas inside of forests, and that way we had a single continuous raster with negative and positive values expressing the distance (km) to the closest forest edge (Morera-Pujol et al. 2022).

Lastly, we obtained the human footprint index, a measure of cumulative human activities, to test whether badgers would be attracted to human structures and habitations. The human footprint index is a unitless index varying from 0 to 50, and it was calculated at a 1km resolution (Venter et al. 2018).

There are environmental variables, such as aspect, that have been shown to have an effect on badger and sett distribution –badgers are more likely to have main setts in south or south-east facing slopes (Byrne et al. 2012), for example. However, if a variable is very granular, that is, it varies very quickly in space (its spatial autocorrelation range is much shorter than the size of the study area), it cannot be included in a large-scale model, as its effect on the badger and sett distribution acts at a much smaller scale. As a result, variables such as aspect, the distribution of small woody features (representing hedgerows), and other habitat types with a much more granular distribution in Ireland.

In addition to the variables shared between the two models, two additional variables were added to the badger model: the output of the setts model was included to examine whether a higher sett density corresponds to a larger number of badgers in the area, and the culling history (badgers culled in the 5 years previous to the data used in the models) was added, to examine whether the persistent perturbance of the culling programme had an effect on badger densities. To produce this variable, a “culled badger density” layer was produced from all the badgers culled between 2013 and 2018 (the 5 years previous to the data in the models) using kernel density estimation (function sf.kde from the package spatialEco, (Evans and Murphy 2025) with a bandwidth of 30 and the same resolution as the covariate rasters). All covariates were scaled before entering the model to ensure better numerical stability and interpretability of the results, and then de-scaled for plotting to aid interpretation.

### Modelling

#### log-Gaussian Cox processes

Setts and badgers were modelled as separate point processes. Spatial point process models are an appropriate technique for modelling presence-only data like the locations of setts or badgers without the need to generate pseudoabsences, and they also have the advantage of being able to account for spatial structure in the data (Renner et al. 2015). To fit these models in a computationally efficient manner, a method has been developed within a Bayesian context that can model point processes as log-Gaussian Cox processes (LGCP), a family of point process driven by a latent Gaussian field, and flexible enough to accommodate ecological processes. Approximating these through an integrated nested Laplace approximation (INLA) method, and using stochastic partial differential equations (SPDE) to approximate the Gaussian field, makes the method flexible and computationally efficient (Illian et al. 2013).

In this context, the data was modelled as an LGCP with the following intensity function:

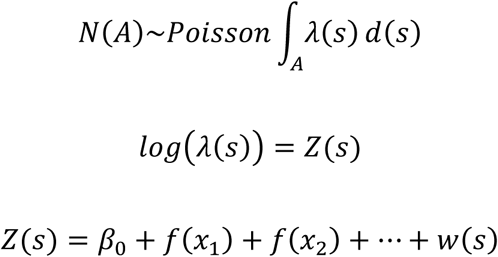

where N is the expected number of points (setts or badgers) in the study area A (in this case Ireland), and λ is the intensity function. The linear predictor Z(s) is connected to the intensity through a logarithmic link and is composed of an intercept (β_0_), several covariates that are modelled with a non-linear function (f(x_i_)) and a spatial random field (w(s)) which will account for spatial autocorrelation and *capture* all the spatial structure in the point data that is not explained by the covariates.

#### The SPDE and the mesh

The complex part of a LGCP is the latent Gaussian Random Field (GRF). To integrate it in a computationally efficient way, an SPDE is used to map the continuous field to a finite set of points defined by a mesh. The mesh is composed of triangles of a size determined by the user, and the triangles vertices (nodes) is where the GRF will be approximated. Between these nodes, the value of the GRF is linearly interpolated (Lindgren et al. 2011).

The definition of the mesh is a crucial step in the modelling process, as there is a trade-off between resolution and computational efficiency: the mesh needs to be fine enough to capture areas where the point intensity changes rapidly, and coarse enough to ensure a reasonable running time. In addition, the covariates will also be evaluated at the location of the mesh nodes, and therefore consideration needs to be given to how well the mesh nodes capture the covariates’ spatial structure. Covariates that are too granular (i.e. vary too rapidly in space) will not be appropriately represented by their values at the mesh nodes unless the mesh is constructed to be very fine, which would make the model computationally unfeasible, and therein lies the limitation of the model in terms of covariate selection (as mentioned above). The code we used to define the mesh in this case is included in the supplementary material for reproducibility purposes.

#### Effort

The integration points that are defined by the mesh nodes also play a part in considering the sampling effort in the model. As mentioned above, the point intensity is evaluated at the nodes and interpolated elsewhere. Prior to running the model each of these nodes is assigned a weight according to the effort, in days, carried out in the quartile where the node falls. Then, when the intensity is evaluated at these nodes, these weights are applied, in a way that means that the model “knows” that some areas have been more intensely sampled than others. Similarly, nodes falling on areas that have not been sampled (e.g. Northern Ireland) will have a weight of 0, which means that the model will not interpret those areas as not containing any observations, but as not sampled at all.

#### Covariate effects

Environmental covariates were assumed to have a non-linear effect on sett and badger distributions. To model this non-linearity we used a smooth function through a 1-dimensional SPDE, which works in a very similar way as the 2D SPDE explained above. The 1-dimensional function is broken into segments (instead of triangles) defined by nodes, and the value of the function is evaluated at the nodes and interpolated between them. The use of priors (explained below) and the number of mesh nodes allow us to control the complexity (wigglyness) of the non-linear effect in a similar way that the number of knots does in a Generalised Additive Model (GAM). More technical details of this interpolation and the role that weights play in accounting for sampling effort can be found in Simpson et al. 2011.

#### Priors

We used Penalised Complexity (PC) priors (Simpson et al. 2014) to set priors on the 1D and 2D SPDE models. PC priors are developed to be easily controlled and interpreted, and to be weakly informative while at the same time avoiding overfitting (penalising complexity). They do so by pulling the field (or function, in the 1D case) to its simplest realisation (i.e. completely flat, with infinite range and a standard deviation of 0). PC priors for the SPDE models are therefore set on the range (ρ) and the standard deviation (σ), in such a way that:

-The prior on the range is set by providing the lower tail quantile ρ0 and the probability P(ρ) of the range distribution so that:

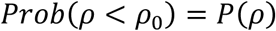

The user defines ρ_0_ and P(ρ) so that, for example, the probability of the posterior range (ρ) being smaller than 5 (ρ_0_ = 5), is 5% (P(ρ) = 0.05), which means we are forcing the range to be between 5 and infinity (flat field).

-The prior on the standard deviation is set by providing the upper tail quantile (σ_0_) and the probability P(σ) of the standard deviation distribution so that:

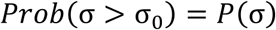

The user defines σ_0_ and P(σ) so that, for example, the probability of the posterior range (σ) being larger than 1 (σ_0_ = 1) is 5% (P(σ) = 0.05), which means that we are forcing the standard deviation to be between 1 and 0 (flat field).

Prior specification is a crucial step of the modelling process, however an established protocol for choosing them does not exist (Morera-Pujol et al. 2022). Following expert recommendations we started with a value of ρ_0_ for the range that was equal to 1/3 the difference between the maximum and minimum value of the covariate (for 1D SPDE models) and 1/3 of the largest dimension of the study area (for the 2D SPDE), and a P(ρ) of 0.5 (i.e. the range is equally likely to be above than below the value of ρ_0_, and for the standard deviation a σ_0_ of 1 and a P(σ) of 0.5). After the first run of the model, we checked the difference between the prior value of ρ_0_ and σ_0_ and their corresponding posterior means and adjusted them accordingly when the posteriors were too far away from the prior values. We iterated this adjustment process until the posterior distributions of ranges and standard deviations looked satisfactory (centred around the mean and without long tails).

#### Model outputs

We used the resulting model to predict sett or badger distribution not only over the sampled area, but over the entire surface of Ireland, including Northern Ireland, projecting it onto a 1km^2^ grid and obtaining a predicted relative distribution of setts or badgers over the island of Ireland. However, although our measure of days spent on the field allowed us to account for spatial differences in effort, there was no way to account for the unobserved setts and badgers, and therefore our estimates were a gross underestimation of the actual sett and badger densities. To avoid confusion, we rescaled those predictions to a measure of relative abundance so the values would go from 0 (lowest badger density) to 1 (highest badger density). Additionally, due to the Bayesian nature of the models we had a measure of uncertainty, as we obtained a distribution of values for every grid cell. The median relative distribution could be then plotted together with the width of the confidence interval, to account for the prediction uncertainty.

Additionally, we evaluated the effect of the covariates at arbitrary values of each covariate (we defined them as 100 values between 1^st^ and 99^th^ quantiles of the dataspace for every covariate).

This produced smooth partial effect plots that we could use to interpret the effect of each covariate.

#### Checking ecological realism of the model predictions using badger weights

Body weight in badgers is thought to be a density-dependent process affected both by sett density and by badger group size in Ireland (Byrne et al. 2015), as well as seasonal effects, so we used Generalised Additive Models (GAM, Wood 2017) through the package *mgcv* (Wood and Sciences 2013) to investigate the effect of sett and badger density, and a proxy for group size (see below), on badger body weight in our study system. From our model outputs we obtained a raster of median sett density and median badger density, and additionally we obtained a proxy for badger group size across the island by dividing the latter by the former. Then, we extracted, at the location of every vaccination badger capture, the values for sett and badger density and for group size proxy, and using the Gamma family (as weight data is 0 bounded) and a log link, we fitted the following model:

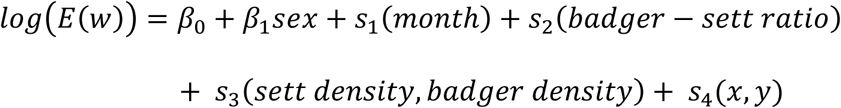

where, with a log link, *E(w)* corresponds to the expected value for badger weight, *β_0_* is the intercept, and *β_1_* the coefficient for a fixed term for the sex of the animal (2-level categorical predictor), as males are usually larger than females. The effect of month (*s_1_*) was modelled to account for seasonal changes in weight, using a cyclic cubic regression spline with a maximum of 10 degrees of freedom (*df*), with the advantage that the cyclic spline forces the model estimates of month 1 to be a continuation of month 12, reflecting the cyclical nature of months of the year. The effect of the badger to sett ratio was modelled as a thin plate regression spline (*s_2_*) with 8 *df*, which models a non-linear relationship. The effects of badger density and sett density were modelled together using a bivariate smooth function (*s_3_*) with 14 *df*, estimated via a thin plate spline assuming isotropy, and spatial autocorrelation was accounted for by including easting and northing (x and y) in another bivariate smooth function (*s_4_*) with 14 *df*.

We ran the INLA models using the functions in the package *inlabru* (Bachl et al. 2019). The package *ggplot2* (Wickham 2009) was used to produce the plots, *sf* (Pebesma 2018) to handle point data, and *gratia* (Simpson 2024) to check GAM model diagnostics. The version of R was 4.2.3 (2023 – 03 – 15 (Team 2023).

## Results

After the filtering process explained in the methods section, the dataset for the sett model consisted of 10,904 set locations (Fig. 1a) last visited between January 2019 and November 2024, whereas the dataset for the badger model consisted of 51,657 badgers (Fig. 1b) in total captured between January 2019 and May 2025. Of those, around 46% were part of the vaccination programme. For the GAM model used to verify the ecological realism of our Bayesian model predictions, we used all badgers captured between 2019 and 2025 within the vaccination programme (before thinning process), and those had a mean weight of 9.68 kg (sd = 1.70) for females and 10.30 kg (sd = 1.76) for males. The effort in number of days per quartile ranged between 6 and 726 days (Fig. 1c).

**Figure 1:**
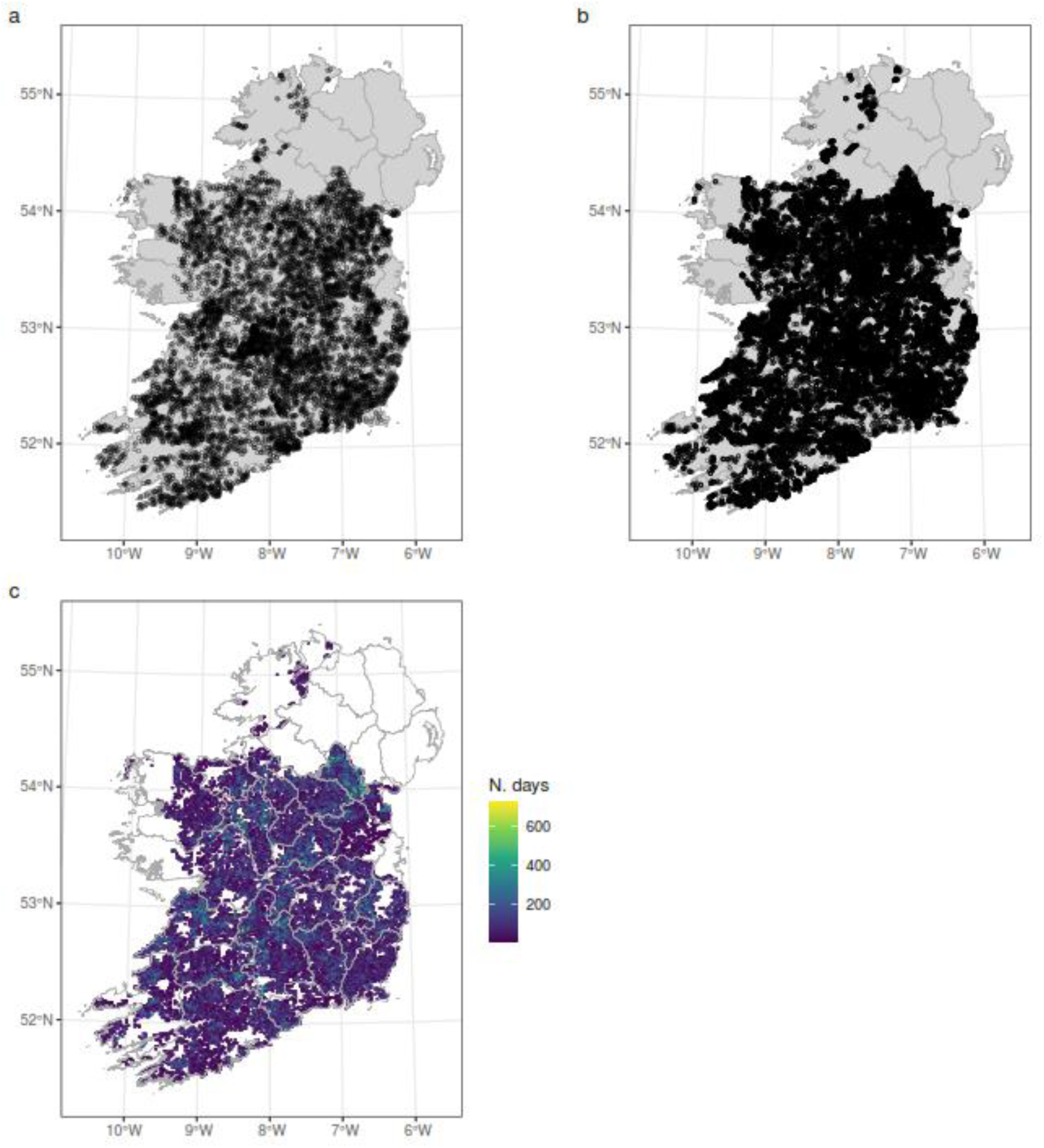
visualisation of the data obtained from the badger culling and vaccination programmes of the bovine Tb eradication programme of the Department of Agriculture, Food, and the Marine from the government of Ireland. Open circles in a) indicate sett the locations of setts last visited between 2019 and 2023. Closed circles in b) indicate the location of badgers captured during the culling programme, or the first capture of the badgers within the vaccination programme. The sampling effort of the culling and vaccination programme combined is shown in c).

### Sett model

We found a clear effect of all variables on the distribution of setts (Table 1), although the effect of human footprint index was the most uncertain (wider credible intervals).

**Table 1:** prior specification (range and standard deviation) and posterior distribution (median and 95% credible intervals) of the covariate effects (1 dimension SPDE) and the spatial random effect (2 dimension SPDE) of the model for the distribution of badger setts in Ireland. Values for the prior and posterior ranges were back transformed with the same parameters used for scaling, so they would make ecological sense.

| Covariate | Prior |  | Posterior (median and 95% credible interval) |  |
| --- | --- | --- | --- | --- |
| Elevation (meters a.s.l.) | $P(\rho) < 414.27 = 0.1$ | $P(\sigma) > 0.1 = 0.1$ | $\rho = 361.34$<br>(256.65 – 571.78) | $\sigma = 0.27$<br>(0.16 – 0.44) |
| Slope (rad) | $P(\rho) < 0.18 = 0.1$ | $P(\sigma) > 0.1 = 0.1$ | $\rho = 0.15$<br>(0.10 – 0.27) | $\sigma = 0.18$<br>(0.10 – 0.30) |
| Grasslands and pastures coverage (%) | $P(\rho) < 60.6 = 0.1$ | $P(\sigma) > 0.1 = 0.1$ | $\rho = 119$<br>(79 – 211) | $\sigma = 0.15$<br>(0.09 – 0.34) |
| Distance to forest edge (meters) | $P(\rho) < 11.93 = 0.1$ | $P(\sigma) > 0.1 = 0.1$ | $\rho = 20.30$<br>(8.25 – 52.94) | $\sigma = 0.16$<br>(0.08 – 0.34) |
| Human footprint index | $P(\rho) < 27.20 = 0.1$ | $P(\sigma) > 0.1 = 0.1$ | $\rho = 23.29$<br>(15.73 – 49.20) | $\sigma = 0.08$<br>(0.04 – 0.17) |
| Spatial effect | $P(\rho) < 100 = 0.01$ | $P(\sigma) = 0.1$ | $\rho = 130.88$<br>(81.05 – 215.27) | $\sigma = 0.1$ |

**Table 2:** prior specification (range and standard deviation) and posterior distribution (median and 95% credible intervals) of the covariate effects (1 dimension SPDE) and the spatial random effect (2 dimension SPDE) of the model for the distribution of badgers in Ireland. Values for the prior and posterior ranges were back transformed with the same parameters used for scaling, so they would make ecological sense.

| Covariate | Prior |  | Posterior (median and 95% credible interval) |  |
| --- | --- | --- | --- | --- |
| Elevation (m a.s.l.) | $P(\rho) < 503.02 = 0.1$ | $P(\sigma) > 0.5 = 0.1$ | $\rho = 879.63$<br>(478.18 – 1,748.85) | $\sigma = 0.73$<br>(0.35 – 1.53) |
| Slope (rad) | $P(\rho) < 0.18 = 0.1$ | $P(\sigma) > 0.5 = 0.1$ | $\rho = 0.16$<br>(0.09 – 0.44) | $\sigma = 0.41$<br>(0.21 – 0.81) |
| Grasslands and pastures coverage (%) | $P(\rho) < 94.11 = 0.1$ | $P(\sigma) > 0.5 = 0.1$ | $\rho = 203$<br>(113 – 518) | $\sigma = 0.28$<br>(0.10 – 0.98) |
| Distance to forest edge (m) | $P(\rho) < 7.07 = 0.1$ | $P(\sigma) > 0.5 = 0.1$ | $\rho = 19.74$<br>(8.23 – 54.99) | $\sigma = 0.18$<br>(0.08 – 0.43) |
| Human footprint index | $P(\rho) < 42.58 = 0.1$ | $P(\sigma) > 1 = 0.1$ | $\rho = 54.88$<br>(26.85 – 138.83) | $\sigma = 0.64$<br>(0.20 – 2.16) |
| Relative sett density | $P(\rho) < 0.64 = 0.1$ | $P(\sigma) > 0.5 = 0.1$ | $\rho = 0.90$<br>(0.53 – 1.96) | $\sigma = 0.26$<br>(0.08 – 0.88) |
| Culling history | $P(\rho) < 0.78 = 0.1$ | $P(\sigma) > 0.8 = 0.1$ | $\rho = 0.99$<br>(0.59 – 1.96) | $\sigma = 0.36$<br>(0.15 – 1.14) |
| Spatial effect | $P(\rho) < 120 = 0.01$ | $P(\sigma) = 0.01$ | $\rho = 79.36$<br>(66.94 – 94.08) | $\sigma = 0.1$ |

We found that lower elevations and shallower (but not completely flat) slopes were more suitable habitat for setts, as well as higher coverage by pastures and grasslands. Sett density was higher inside or near the edge of forests, and although the effect of the human footprint index was more uncertain, there seemed to be more setts in areas with moderately high human activity. (Fig. 2).

**Figure 2:**
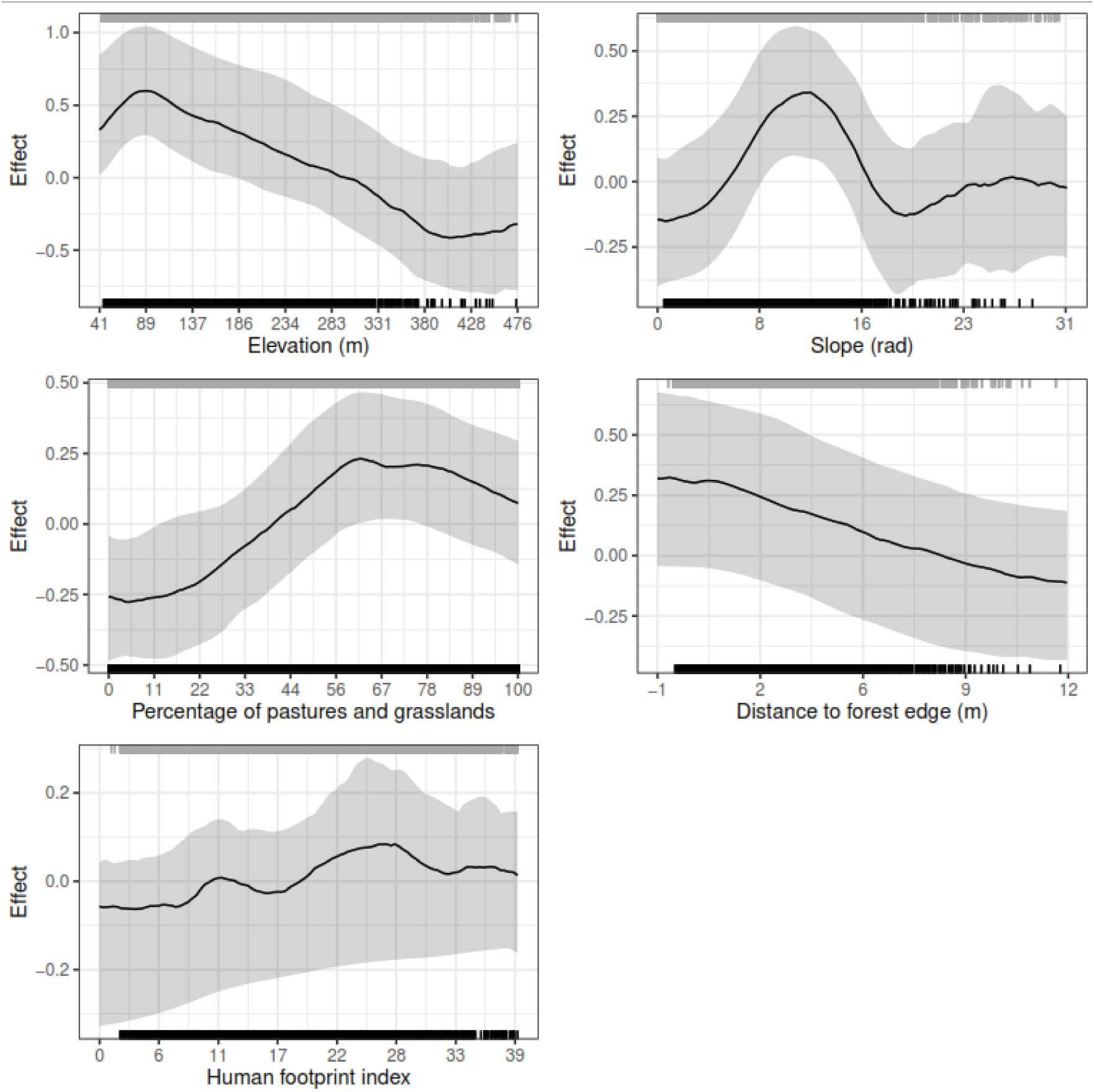
non-spatial evaluation of the covariate effects of the sett distribution model at 100 values evenly distributed from the minimum covariate value in the study area to the value of the 99^th^ percentile (to avoid extreme values). The grey rug on the top shows the covariate values at the integration points (distributed evenly across the study area) and the black rug at the bottom shows the values at the observations. The solid black line shows the median covariate effect values, and the shaded ribbon shows the 95% credible interval obtained from drawing 100 samples from the model. The values in the x axis have been back transformed using the scaling parameters so they would be easier to interpret.

Predicting from the sett model, we obtained a spatially explicit posterior distribution for badger sett abundance for the whole island (including Northern Ireland and the unsampled areas, Fig. 3a). We plotted the median and 95% credible interval width (as a measure of uncertainty) of that distribution to show the lowest densities of badger setts in the mountainous areas in the Wicklow mountains (to the East), County Kerry (in the Southwest) and Connemara (West), and the highest densities in the Southeast (coasts of counties Wexford, Waterford and Cork), with intermediate sett densities elsewhere. This distribution is mainly driven by elevation, with the other covariates contributing as well but with a smaller effect size (Fig. S1). The uncertainty is higher in mountainous areas as it is to be expected due to the fact that little to no observations exist in those areas (Fig. 3b).

**Figure 3:**
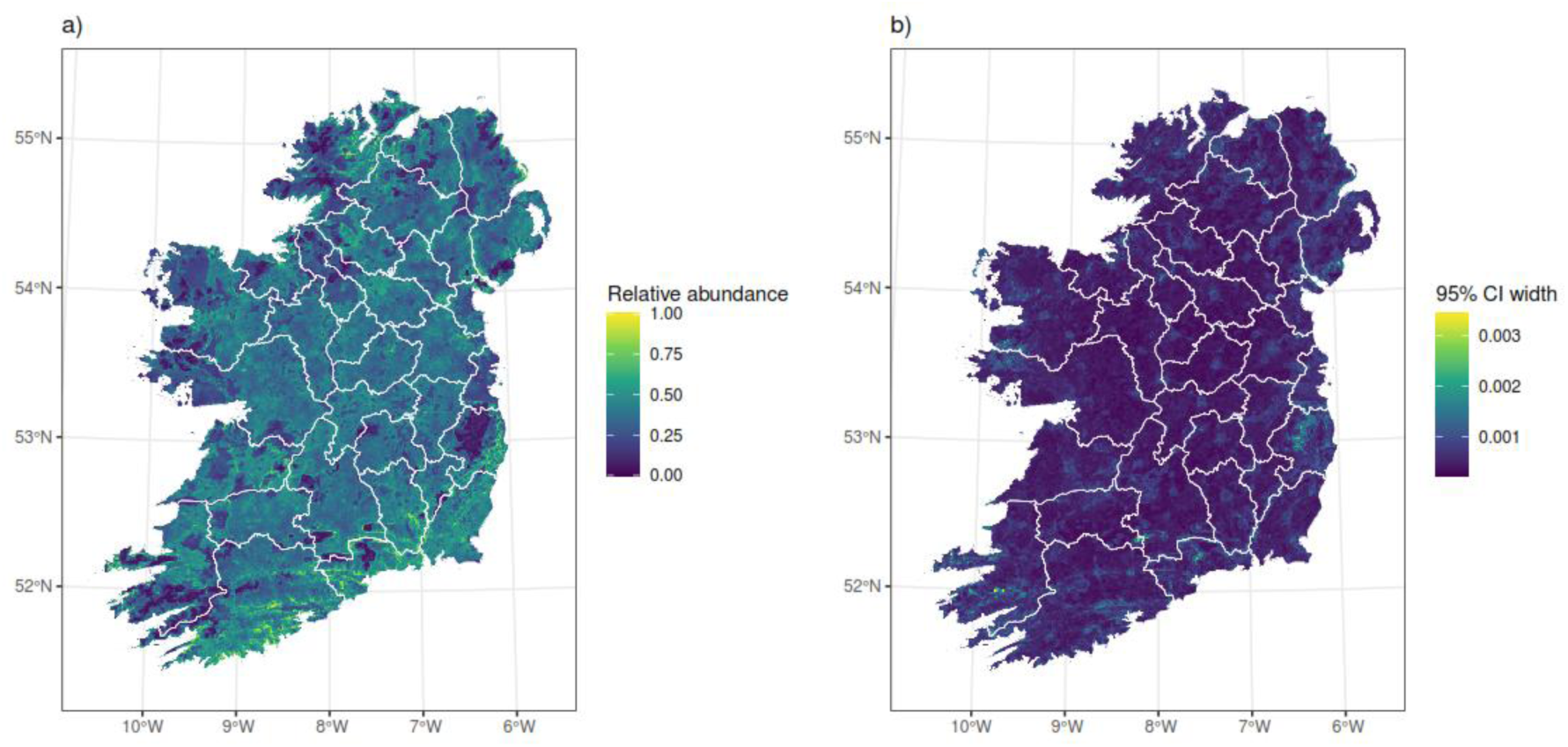
badger sett predicted distribution obtained from the spatially explicit model on a 1×1km grid. The Bayesian nature of the model produces a posterior distribution of predicted sett density for every grid cell, which allows us to plot the median sett density (a) and the credible interval width (b) reflecting the uncertainty of the model.

The spatial random field captured a gradient (albeit of a very small magnitude) with higher sett density to the Southeast of the country, that could not be explained by the environmental covariates in the model (Fig. S2).

### Badger model

We found that all environmental variables except human footprint index had an effect on the distribution of badgers. Of the additional, badger-specific variables, culling history had a much stronger effect than relative sett density.

We found that badgers showed a preference for lower and medium slopes and elevations, and a clear –although smaller in magnitude– preference for higher coverage of grasslands and pastures and shorter distances to forests. Human footprint index did not seem to have an effect on badger density, and neither did the relative abundance of setts. Finally, we found that areas where badgers have historically been abundantly culled have lower badger densities (Fig. 4).

**Figure 4:**
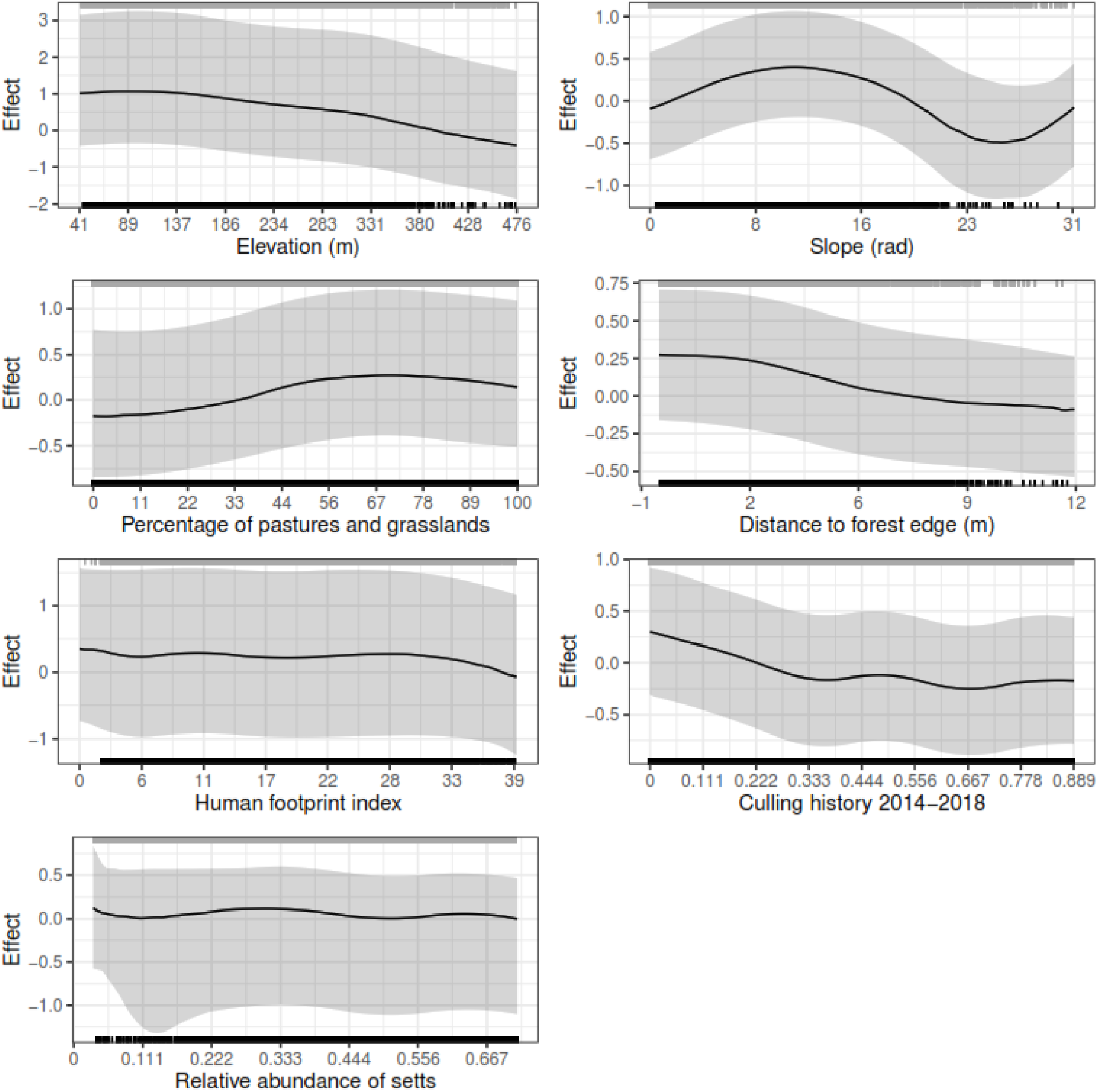
non-spatial evaluation of the covariate effects of the badger distribution model at 100 values evenly distributed from the minimum covariate value in the study area to the value of the 99^th^ percentile (to avoid extreme values). The grey rug on the top shows the covariate values at the integration points (distributed evenly across the study area) and the black rug at the bottom shows the values at the observations. The solid black line shows the median covariate effect values, and the shaded ribbon shows the 95% credible interval obtained from drawing 100 samples from the model. The values in the x axis have been back transformed using the scaling parameters so they would make ecological sense.

**Figure 5:**
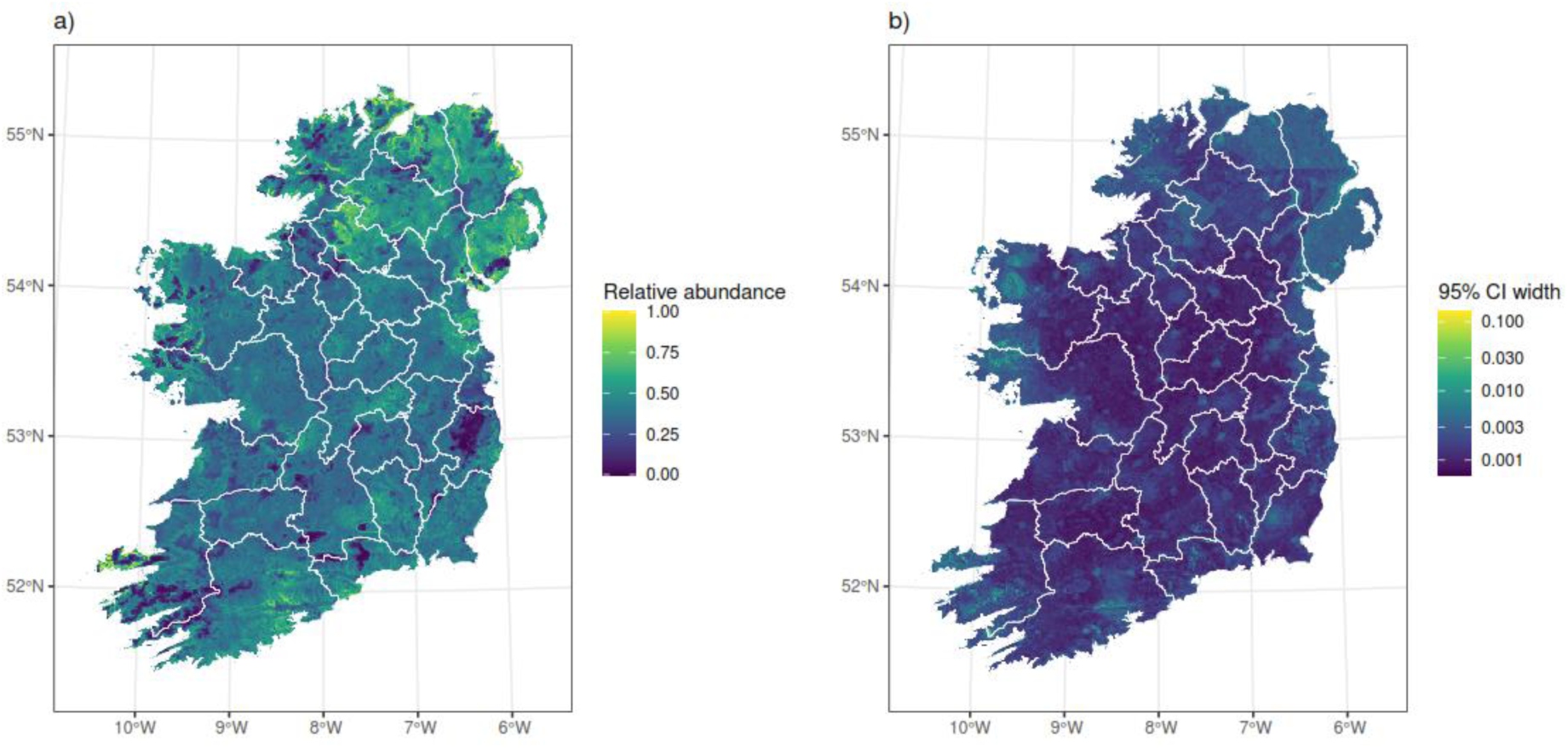
badger predicted distribution obtained from the spatially explicit model on a 1×1km grid. The Bayesian nature of the model produces a posterior distribution of predicted badger density for every grid cell, which allows us to plot the median sett density (a) and the credible interval width (b) reflecting the uncertainty of the model.

The median of the spatially explicit prediction showed low badger densities in the mountainous areas in Wicklow, Kerry and Galway, and higher badger density in Northern Ireland and some regions of the southeast. These distributions are mainly driven by elevation with culling history (i.e. the lack of culling) being responsible for the higher densities in Northern Ireland (Fig. S3).

Uncertainty was more or less uniform across the study area, except for some areas in the mountains where there were less observations and therefore uncertainty is higher, and Northern Ireland where the lack of data had the same effect.

The spatial random field shows higher densities in the East and Northwest unexplained by the model covariates (Fig. S4)

### Ecological realism of model predictions

The GAM model revealed significant effects of all terms (with significance level set at p = 0.05), and explained 34.9% of the deviance. Model diagnostics revealed no relevant structures in the residuals (Fig. S5), and the variogram of the residuals showed no spatial structure (Fig. S6). Males were heavier than females, with an estimated coefficient of 0.06 (SE = 0.002). All smooth terms were significant, with heavier badgers in October and November and lighter in early summer (before and after winter torpor, respectively). Spatially, badgers seemed to be lighter in the North, and of higher body weight in the midlands and Southeast of the country (Fig. S7).

Badger and sett density together had a significant effect on badger body weight where larger badgers were found in areas with more setts and more badgers, and lighter badgers were found in areas with less setts and less badgers. However, group size proxy – that is, the number of badgers per sett per km^2^ – had a negative, almost linear relationship with body weight, where weight decreases with group size (Fig. 6).

**Figure 6:**
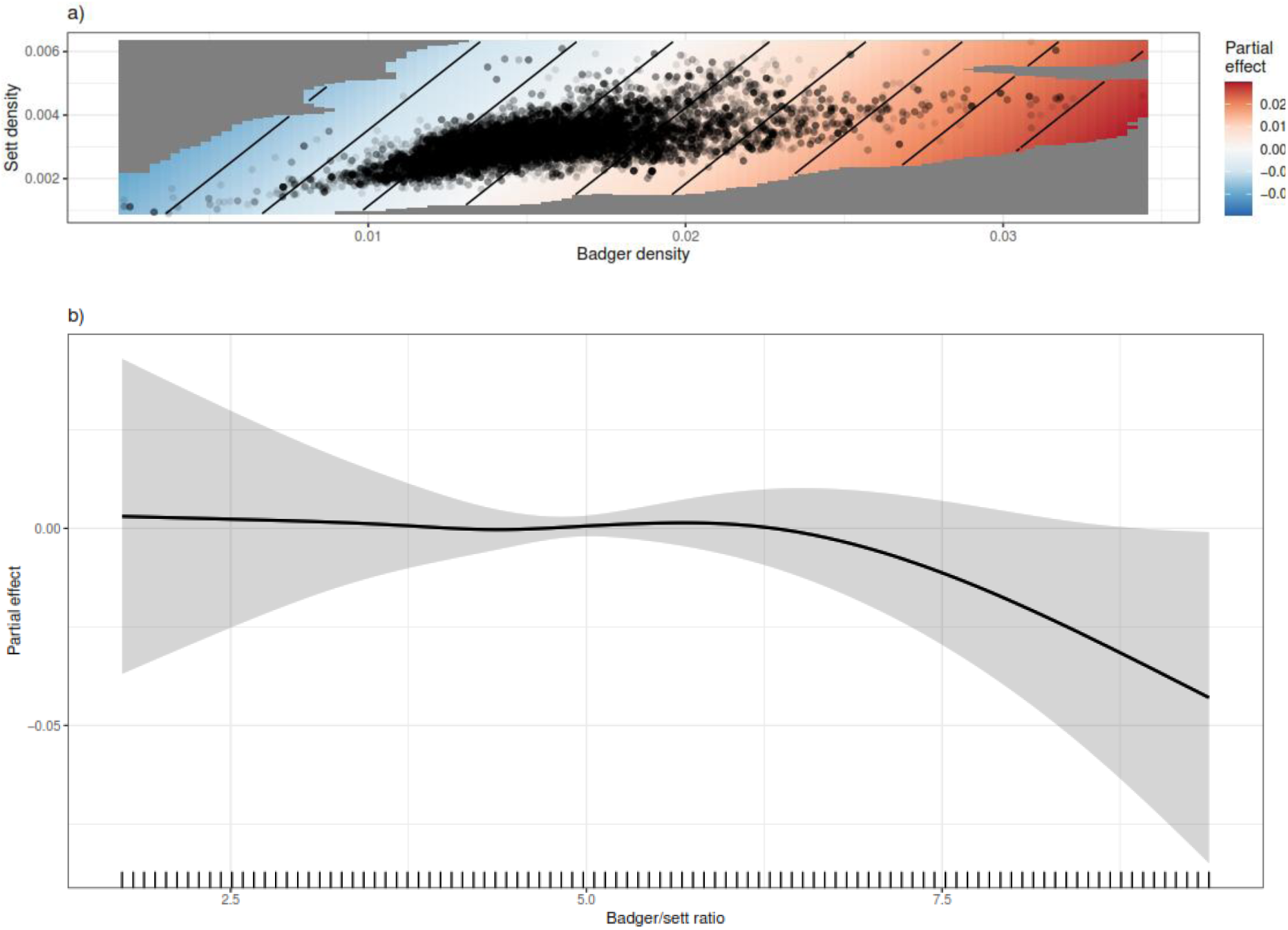
main results of the GAM model. a) shows the effect of badger (x axis) and sett density (y axis) on badger weight when modelled as a bivariate thin plate spline assuming isotropy. Red colour represents heavier badgers, and blue colour lighter badgers. The size of the axes shows the different range in values between sett density (y axis) and badger density (x axis). b) shows the relationship between the group size proxy and weight. Badger weight estimates decrease with an increase of group size proxy (badgers per sett per km^2^)

## Discussion

We showed with this study, for the first time, that the distribution of badger main setts does not necessarily reflect the distribution of badgers themselves, especially in post-cull landscapes, but rather these are two different ecological processes with different environmental drivers. Despite our models not producing estimates of total abundance of setts or badgers, the relative abundance surfaces obtained allowed us to show the decoupling of the two processes. This has relevant consequences for management, as main setts have been commonly used to represent badger groups, and then multiplied by an estimated group size to obtain estimates of badger abundance (Reid et al. 2012, Byrne et al. 2014). Such simple indices of abundance are particularly problematic where populations are culled, vaccinated or both (Byrne et al. 2021), leading to differing population growth trajectories relative to habitat carrying capacity. The complex relationship between sett density and badger density is well captured by our models and reflected in their ecologically meaningful relationship to badger weight.

### Ecological drivers of sett and badger distribution

Badger populations organise in groups or clans, each of them loosely defending a territory that typically contains a main sett and several secondary setts (Byrne et al. 2012). The distribution and size of territories (and therefore the main sett density) depend on habitat, with higher quality habitats able to support more clans as they require smaller territories (Feore and Montgomery 2006, Byrne et al. 2015). Our models showed that flat, low-elevation lands are able to support higher sett densities, together with areas with partial grassland and pasture cover. Forest cover has previously been identified as a driver of sett density (Revilla et al. 2006, Piza Roca et al. 2014) with badgers preferring to construct setts under forest cover and particularly at the edge of forests with other environments like pastures that might provide more abundant food sources. We constructed our covariate specifically to detect the preference for forest edges (as a continuous variable “distance to the forest edge” with negative values for locations inside forests and positive values outside). However, the nature of the sampling design (mainly concentrated around farms) and the increased difficulty in finding sett entrances within forests might have biased these results: although we did not detect a specific preference for the forest edge itself, we did detect a reduction in sett density with increased distance from forests, suggesting that in Ireland badgers prefer to make their setts in or close to forests. It is known that in areas with low forest cover badgers replace the shelter of trees by that of hedgerows (O’Brien et al. 2016), which, together with the added food sources that pastures and grasslands provide (Cleary et al. 2009) might explain their preference for areas with partial pasture cover as hedgerows mark the edges of fields and pastures. We also found a weak but positive effect of human footprint index that dropped at very high values. In a country like Ireland, with farming encroaching in most of its habitats, this might just be reflecting the association with hedgerows and pastures already described, and the negative effect at very high values of human footprint index is likely reflecting an avoidance of built-up areas like towns and cities. More granular features that might also govern the distribution of setts at a local scale, like soil type and slope aspect could not be accounted for in this model at a national scale.

Overall, badger density was driven by similar environmental variables as sett density, being more abundant at low elevations (but with a less pronounced effect) and medium slopes, and in areas with high coverage by pastures and grasslands inside or close to forests. Surprisingly, sett density did not seem to have an effect on badger density, although our GAM model did show that areas where our models predicted higher sett density, they also predicted higher badger density (see below). This correlation might have been masked by the environmental covariate effects, that reflected sett and badger density similarly. Lastly, we included in the model a layer representing culling history. This was calculated by obtaining the kernel density estimation of all badgers culled in the five years previous to our study, and was aimed at capturing the effect that consistent historical removals might have had in the badger population (Byrne et al. 2013b). We found that indeed, areas that had been more intensely culled in the five years previous to the start of the study had lower badger densities, up to a point, and then increased number of badgers historically culled did not have a decreasing effect in current badger density. This might hint to the fact that after intense removal, density dependent effects start to disappear as the population is far away from carrying capacity.

When we modelled the weights of the vaccinated badgers against the sett and badger density and the group size proxy (ratio of badger to sett relative abundances) to test the realism of our model predictions, we found results that overall matched our expectations based on published data. Most badgers live in areas with intermediate sett and badger densities, however the few badgers that live in areas with high sett and badger densities are heavier, and those that live in areas with either low sett or low badger density (or both) are of lower body weight. Therefore, areas with better habitats and therefore more resources will be able to sustain more setts, and more, healthier badgers, while areas with poorer habitats will sustain less badgers and in a poorer body condition. In agreement with (Macdonald et al. 2002), we found a negative, but small, relationship between group size and body weight. This relationship might not be as strong in Ireland as in other populations due to the continued historical removal of individuals through the culling programme, that might have left Irish badger populations well below the ecosystem’s carrying capacity, and therefore density-dependent effects on weight are not as prevalent.

### Badger and sett distribution

Our models produced distribution maps that represented relative abundances (rescaled to a 0-1 gradient), as the incorporation of weights to the integration points could account for different sampling efforts, but it was not a true observational process that would have allowed us to account for unsampled setts. However, they are a good and valuable approximation of relative density for both setts and badgers and show that, as expected, places with low density of setts will have low density of badgers (as no badgers live where there are no setts), but at higher sett densities, however, the two distributions are decoupled. While higher densities of setts are found to the Southeast, which is likely driven by more favourable climate and landscape quality (Byrne et al. 2015), higher densities of badgers are found in Northern Ireland, likely related to the lack of historical culling, as widespread culling has never been authorised, and the only badger removal was in one small area (100 km^2^) Test Vaccinate and Remove (TVR) study in County Down (O’Hagan et al. 2021) with low number of removals (n=108).

### Implications for bTB management

Although the control of cattle-to-cattle transmission and residual herd bTB levels are key aspects of bTB eradication in Ireland (More 2024), badger-to-cattle transmission does occur (Griffin et al. 2005, Chang et al. 2023), and requires careful management. The Government of Ireland invests heavily in its bTB eradication programme (Ryan et al. 2023) and therefore there is growing interest for knowledge that can help design more targeted, cost-efficient actions. Knowing how setts are distributed spatially, and where the higher densities are, is important for management as sett density has implications for bTB spread: badgers living at low sett densities (with larger territories) seem to be less territorial and travel further away from the natal sett, with the possibility to spread bTB to areas previously uninfected (Rogers et al. 1998, Frantz et al. 2010, Byrne et al. 2019).

Group size itself (as opposed to territory size) has failed to be directly linked to bTB prevalence amongst badgers (Vicente et al. 2007, Konzen et al. 2024), although it has been suggested that larger group sizes could facilitate intra-group transmission, which could have an effect on bTB prevalence in the overall badger population (Hardstaff et al. 2012). Additionally, it has been shown that badgers living in smaller groups are also more mobile (Byrne et al. 2022), which might again be related to bTB spread. The data collection method used for this study does not allow for an estimate of group size, however we obtained a proxy for it by dividing the estimated badger density surface by the sett density surface (before rescaling them to a 0-1 scale), and therefore obtaining a measure of “badgers per sett per km^2”^, which can be understood as a proxy for group size (Fig. 7). This has shown larger estimated group sizes in Northern Ireland (probably related to higher badger densities there due to lack of culling), and in the inhospitable areas in the west coast of Ireland, which are probably a result of the badger model somewhat overestimating badger densities in the region, since we know that it was more sparsely sampled, the habitat in that region is not favourable for badgers and there are very few badger setts (Byrne et al. 2014) and smaller group sizes (Feore and Montgomery 2006, Judge et al. 2017). However, the remaining hotspots seem to coincide with areas where the last national bovine TB statistics of the Department of Agriculture, Food, and the Marine report higher prevalences of bTB in cattle herds (Department of Agriculture 2025). Although this may be the result of a more intense capture effort in areas that experience recurrent bTB breakouts, the relationship between badger group size and bTB spread merits further investigation.

**Figure 7:**
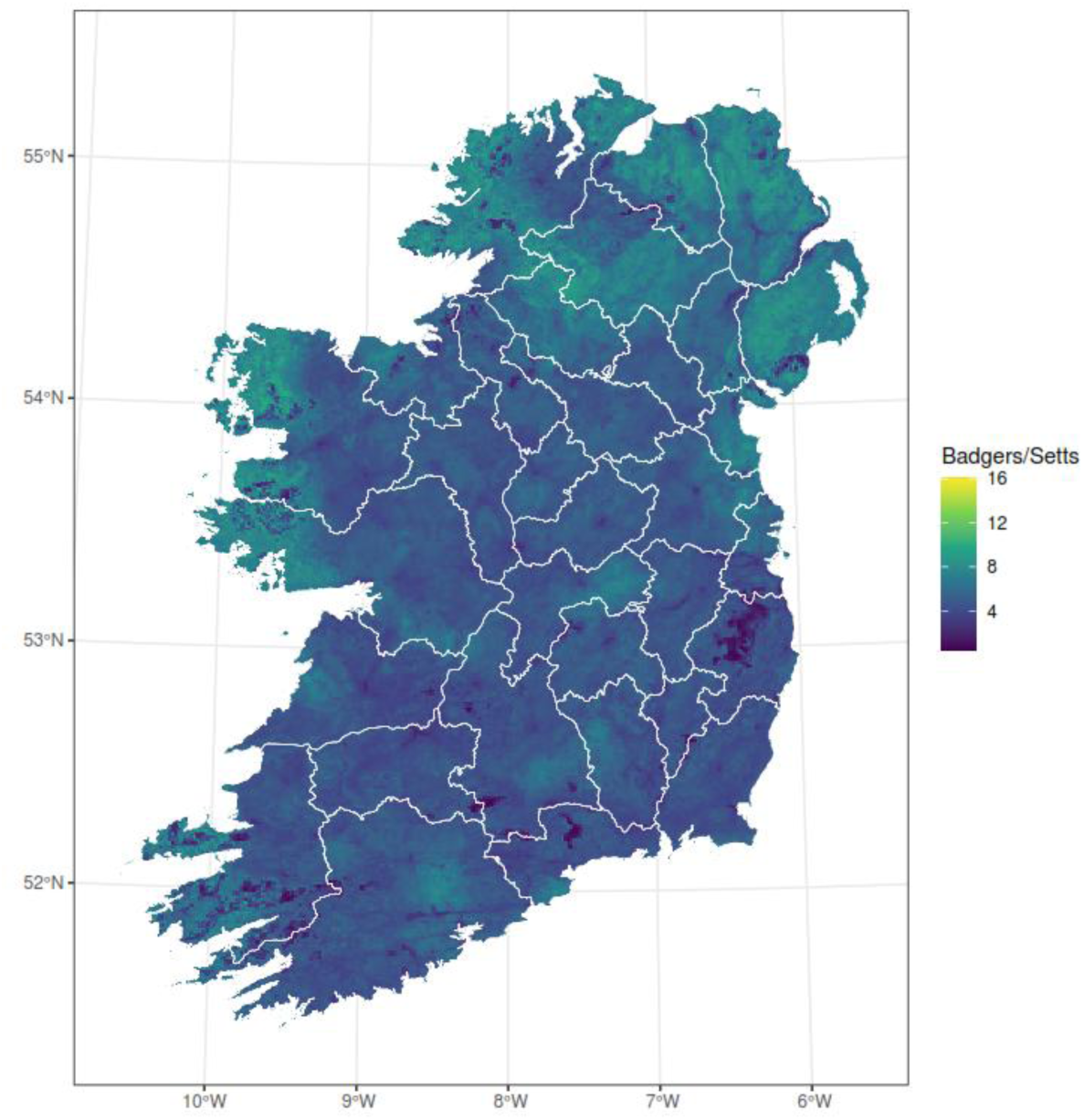
ratio of badgers to setts per 1km^2^, obtained from dividing the badger distribution by the sett distribution. This can be used as a proxy for badger group size.

### Effects of management on badger populations

Badger populations in Ireland have been influenced by management plans since at least 2004. Originally, a culling programme that aimed at reducing the badger density in bTB problematic areas by 25-30% (O’Keeffe 2006), moving on to a more targeted culling plan, and finally by enforcing a replacement of culling with vaccination, a plan that is still currently being rolled out (Byrne et al. 2024). All these management actions have had an impact on the population size and dynamics that remains to be studied, although some efforts have been made to study their effect on movement (O’Hagan et al. 2021, Redpath et al. 2023). Our results show how areas where culling has happened more intensely in the past have a lower density of badgers, matching the results of previous studies that concluded that the culling regime had reduced the badger populations, particularly in favourable environments (Byrne et al. 2021).

As of 2021, more than half the surface of Ireland on which the wildlife unit of DAFM has been designated for vaccination, where the goal is to cull only in response to severe epidemiological cases (Ryan et al. 2023). This change in management has meant that areas under the culling programme in the past, that had been enduring constant removals, are now under the vaccination programme and not experiencing removals anymore, which might cause population numbers to rapidly increase (in a process akin to that of predator release when a top predator is removed from an ecosystem). Additionally, as the roll-out of the vaccination programme progresses, the badger population is expected to be healthier, with fewer diseased animals, which might have consequences in population dynamics and survival (Robinson et al. 2012).

Density dependent population parameters like recruitment or migration will also be affected by these changes (McDonald et al. 2016), which complicates the estimation of population sizes, which is already a complex endeavour (Byrne et al. 2021). Other parameters such as the number of occupied non-main setts per territory (Rogers et al. 2003), or the distances travelled during ranging movements, might also be affected by badger density and therefore the different management regimes, and will merit further investigation as they have relevance in the spread of bTB to cattle.

## Conclusion

Decades of management in Ireland have altered the badger population to a point where estimating badger abundances, densities, and distributions is a complicated endeavour, and estimating population parameters such as mortality, migration, or recruitment is nigh impossible with the currently available data. However, a better knowledge of the badger population in Ireland is essential for an effective control of spillover from badgers to cattle, and therefore for the bTB eradication programme. Our work provides an avenue of future research as it presents a clear image of the decoupling of sett and badger abundance and the importance of obtaining accurate estimates of both.

## Supplementary figures

**Figure S1:**
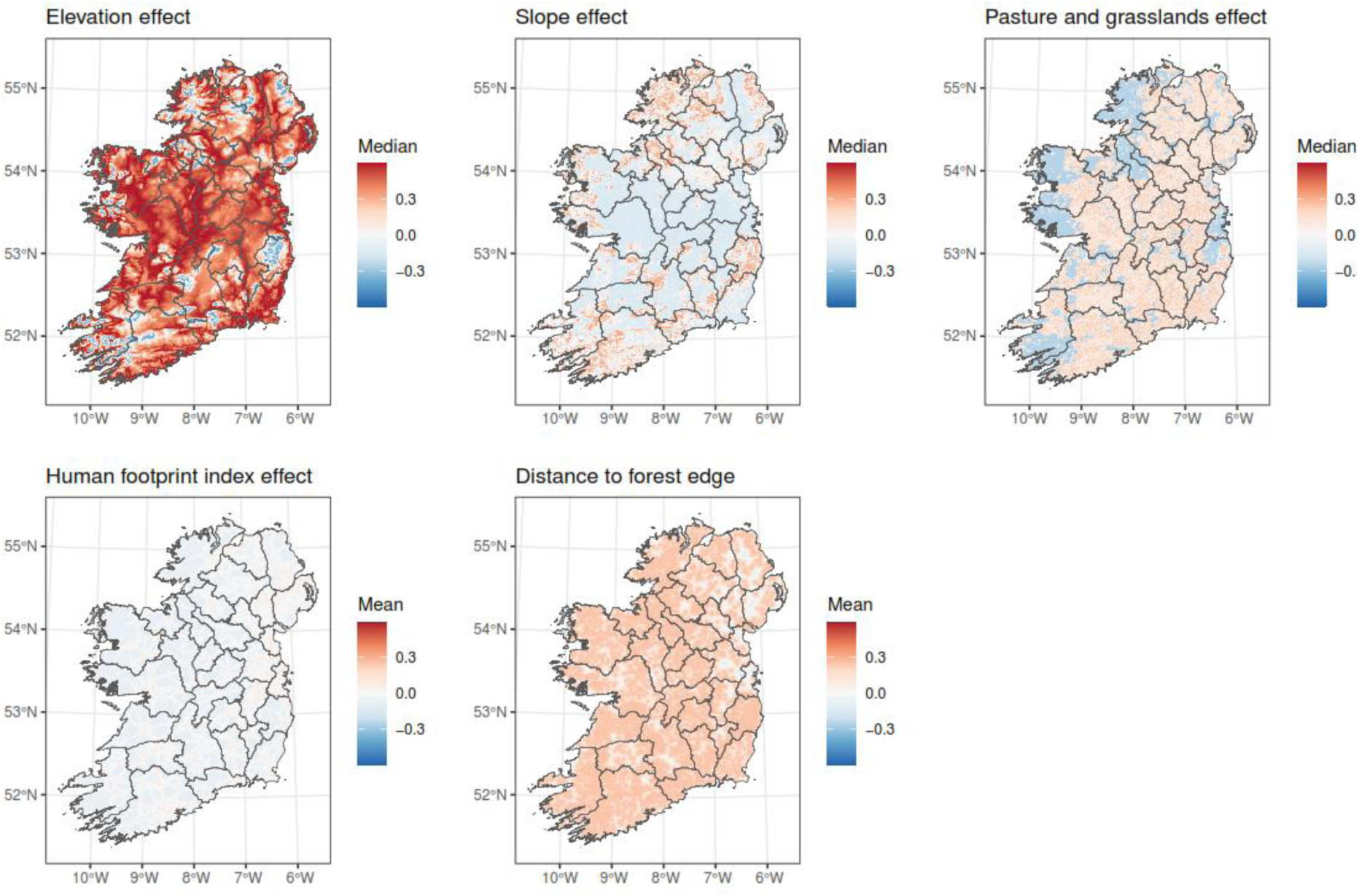
spatially explicit effect of each of the covariates on the model of sett distribution

**Figure S2:**
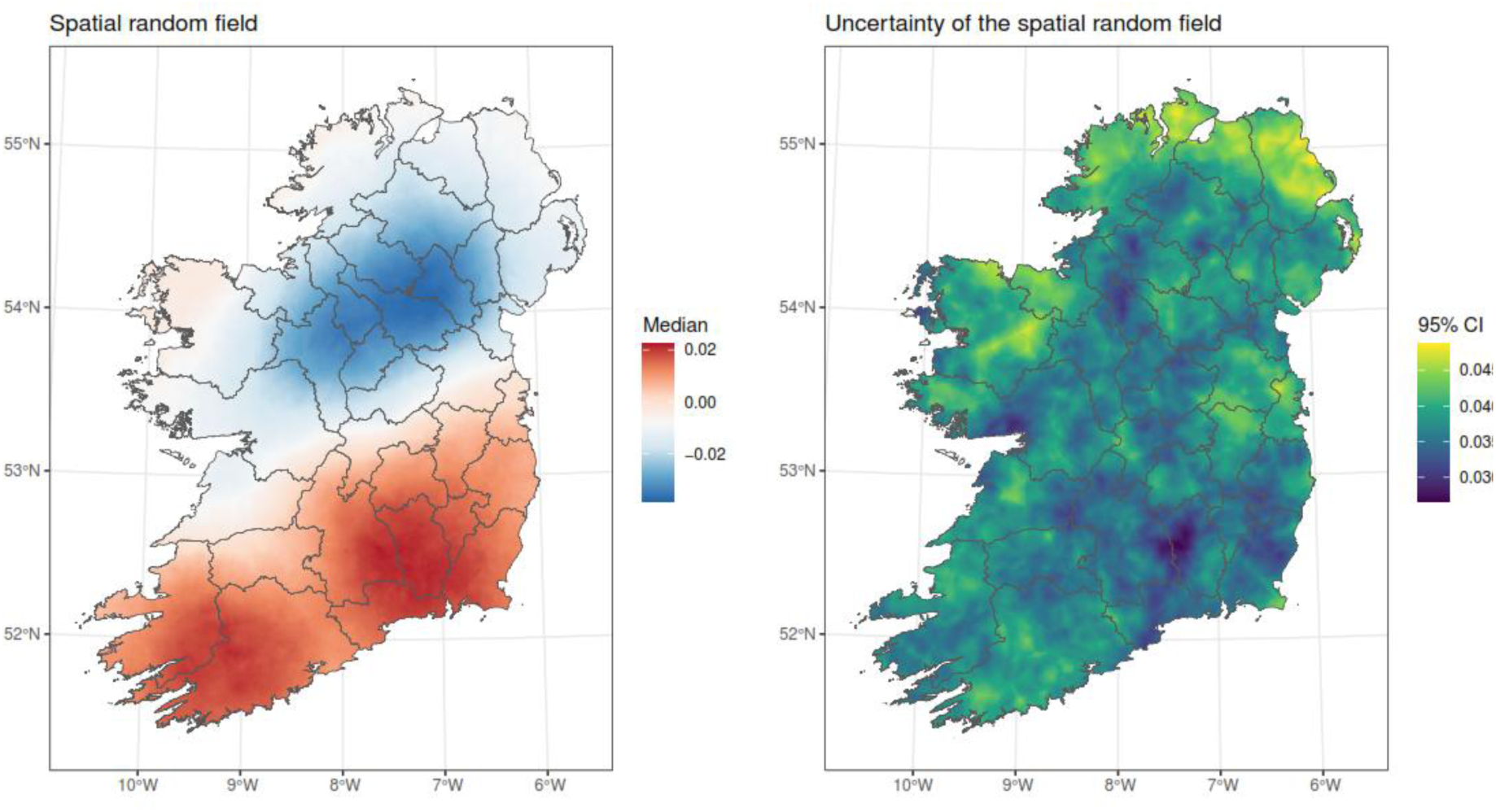
spatially structured random field (left) and its 95% confidence interval, capturing all unexplained spatial structure in the model of sett distribution

**Figure S3:**
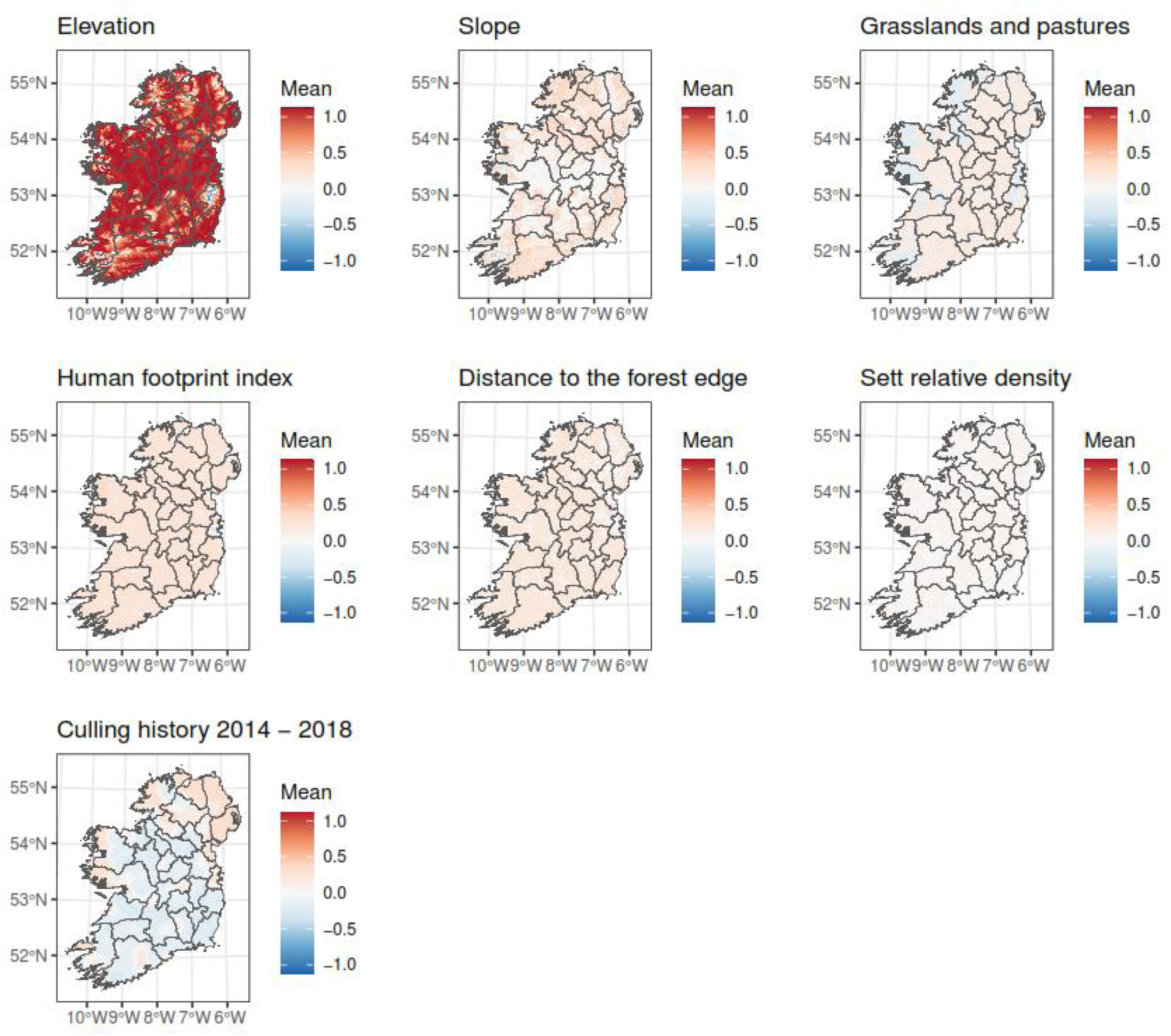
spatially explicit effect of each of the covariates on the model of sett distribution

**Figure S4:**
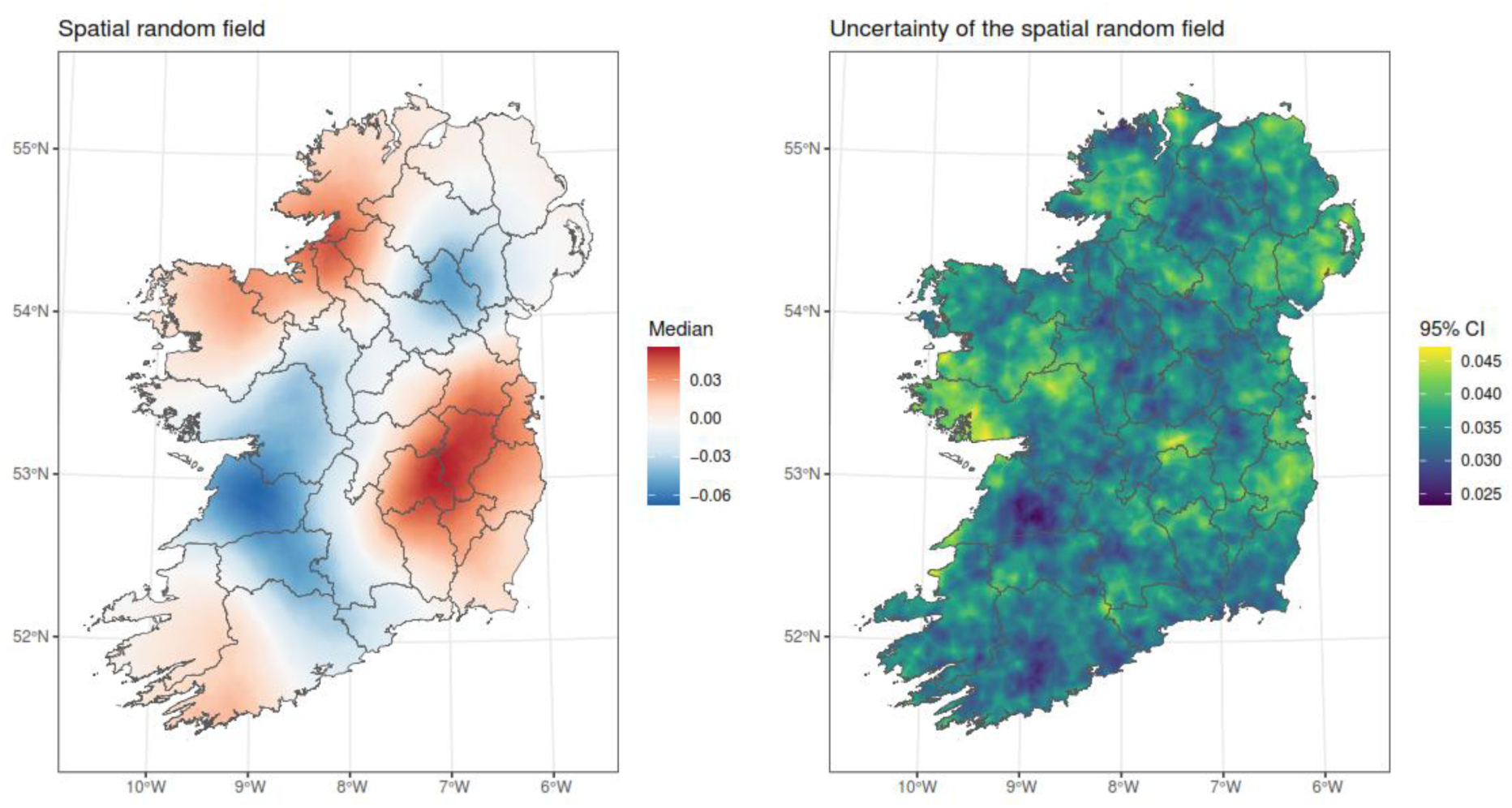
spatially structured random field (left) and its 95% confidence interval, capturing all unexplained spatial structure in the model of badger distribution

**Figure S5:**
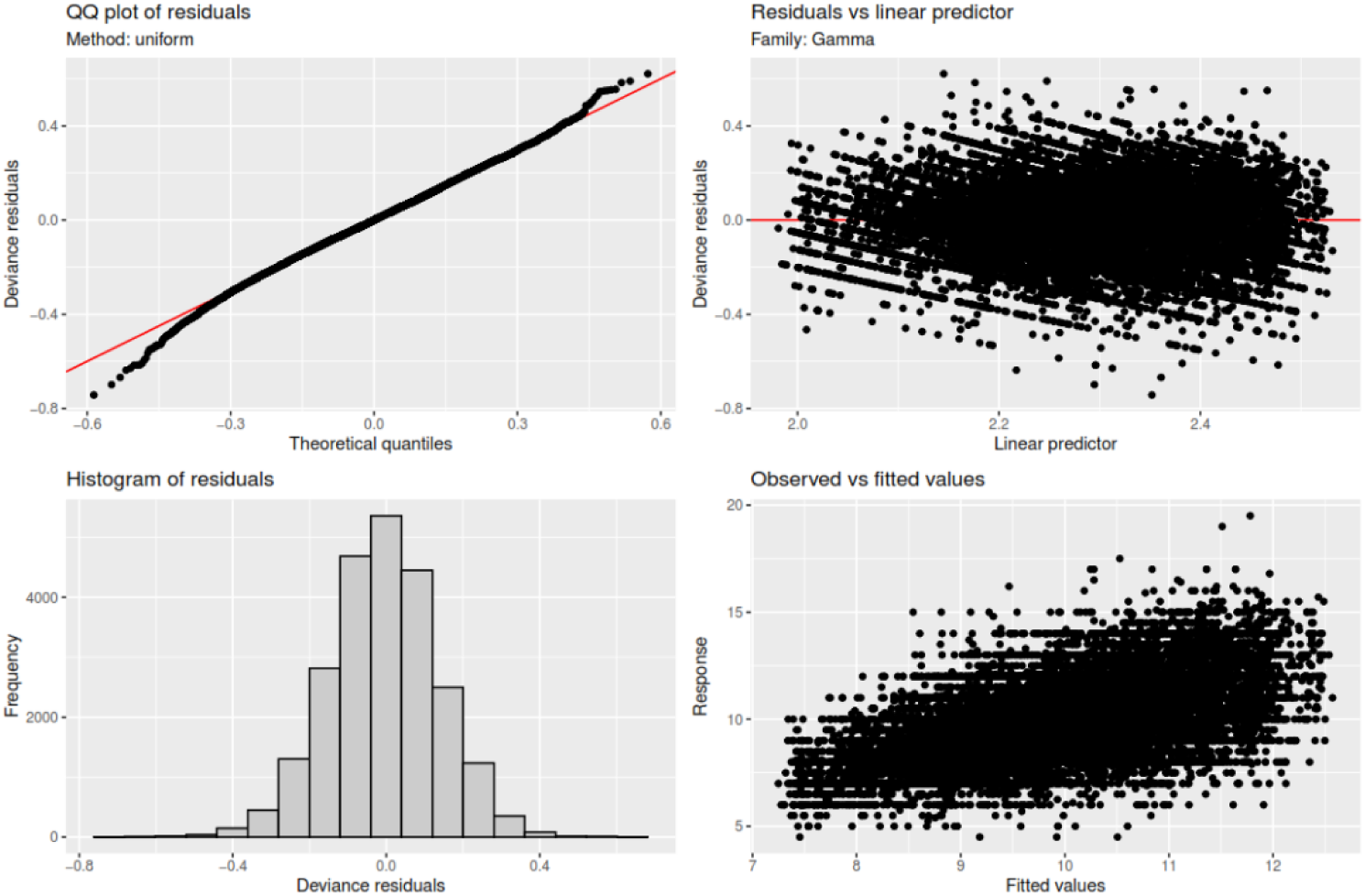
diagnostic plots for the GAM model fit to the badger weight data with sex, group size proxy, sett and badger density and x and y coordinates as explanatory variables. We consider these diagnostics satisfactory.

**Figure S6:**
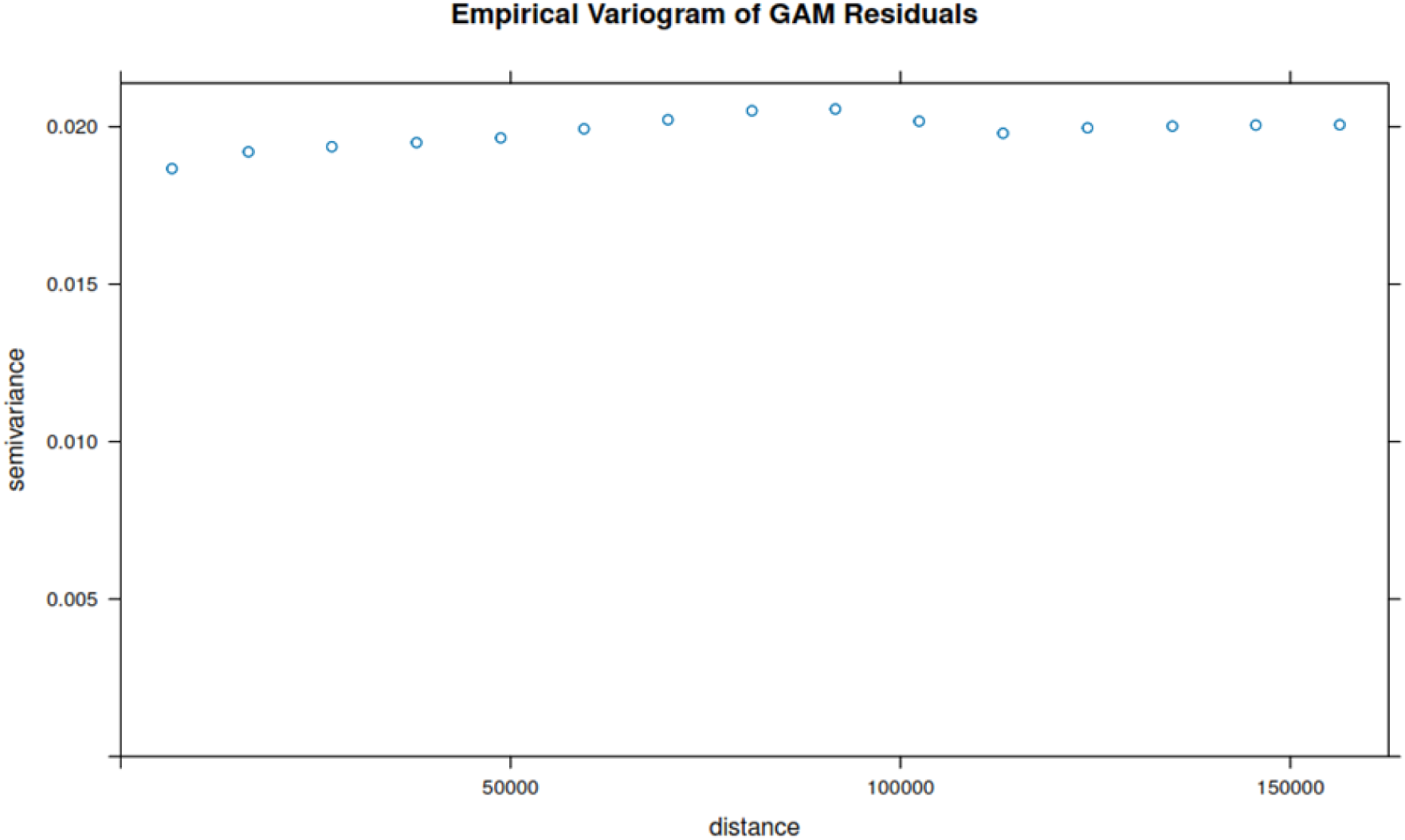
semivariogram of residuals of the GAM model model fit to the badger weight data with sex, group size proxy, sett and badger density and x and y coordinates as explanatory variables, showing no spatial structure of the residuals

**Figure S7:**
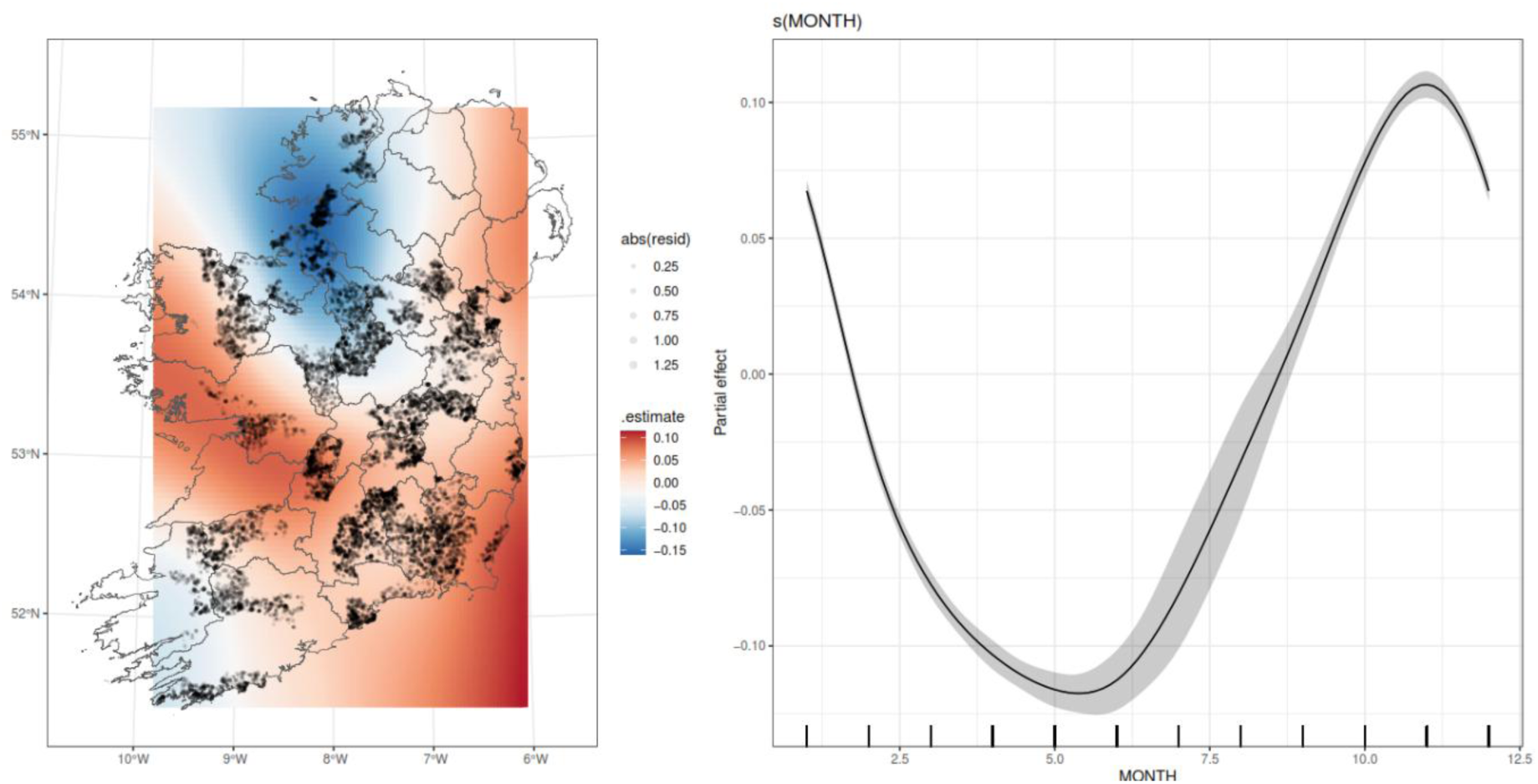
additional results from the GAM model. Left plot shows the effect of longitude and latitude on weight (with red colours delineating areas with heavier badgers and blue colours areas with thinner badgers), with the size of the residual mapped to the size of the points to show the lack of spatial structure on the residuals. Right panel shows the effect of month on badger weight, with a cyclical nature.

